# The effect of chronic, latent *Toxoplasma gondii* infection on human behavior: Testing the parasite manipulation hypothesis in humans

**DOI:** 10.64898/2026.03.16.712071

**Authors:** Kim Valenta, Nicholas Grebe, Thomas Kelly, Jennifer W. Applebaum, Adam Stern, Jordan Traff, Siddharth Satishchandran, Stacy Rosenbaum, Valerie Lantigua, Alice C. Y. Lee

**Affiliations:** University of Florida, Department of Anthropology; Department of Psychology, Occidental College; Department of Environmental and Global Health, University of Florida; Department of Comparative, Diagnostic, and Population Medicine, College of Veterinary Medicine, University of Florida; Department of Anthropology, University of Michigan

## Abstract

Parasitism is one of the key, structural, interspecific interactions in ecology. One remarkable parasitic strategy that has been documented in multiple systems is the behavioral manipulation of hosts to increase parasite fitness. While not yet documented in humans, we propose that a ubiquitous zoonotic parasite – *Toxoplasma gondii* – may change human behavior to favor the parasite by increasing the fitness of the parasite’s definitive host - cats. Specifically, we assess the possibility that human behavioral changes resulting from chronic, latent *T. gondii* infection lead to measurable changes in attitudes, actions and dopaminergic responses towards cats that function to increase domestic cat fitness. We assessed the potential role of humans in the *T. gondii* lifecycle by identifying and testing behavioral changes in humans that benefit the parasite; specifically, human affection for cats. We assessed *T. gondii* infection status in 68 participants using *T. gondii* serum antibody testing, and assessed their attitudes towards cats in three ways: i) surveys, ii) participant behavior in the presence of domestic cats, and iii) participant oxytocin levels before and after interactions with cats to assess dopaminergic changes. Only 2 of 68 participants were positive for *T. gondii* antibodies, limiting statistical power. However, our results indicated that *T. gondii*-positive participants both reported a greater affection for cats in surveys, and spent more time engaged with cats during behavioral trials than *T. gondii*-negative participants (87% of study time engaging with cats vs 75%). Oxytocin results were inconclusive.

## Introduction

Along with mutualism, competition and predation, parasitism is one of the key, structural, interspecific interactions in ecology (1). Parasitism refers to any relationship between two or more species in which one species (the parasite) garners a fitness benefit from a given interspecific association, while the other (the host) incurs a cost. Host-parasite interactions drive variation at several scales, from regulating host demography at the population level, to individual effects on hosts, and host competitors (2). Parasitism is not only an important feature of biotic communities, it is also ubiquitous, with parasites accounting for approximately half of extant biodiversity (3). The diversity of parasites may partly explain the remarkable diversity of parasitic survival strategies, from the obvious, e.g., directly consuming host tissue for nutrients, to the astonishing, e.g., castrating hosts to redirect energy to parasite growth (4).

One particularly remarkable strategy of parasites is the behavioral manipulation of hosts to increase parasite fitness (5). While the idea of a parasite manipulating host behavior for its own purposes is extraordinary, it is not new – behavioral manipulation of hosts by parasites has been suspected for nearly 100 years (6, 7), and has been documented in multiple systems. Laboratory and field studies demonstrate that behavioral manipulation by parasites is utilized for several purposes, including increasing parasite transmission to new hosts, dispersal to favorable environments, and protection from predators (5, 8, 9). For example, neotropical ichneumonid wasps parasitize spiders and manipulate them into weaving webs to protect developing wasp larvae (10). Horsehair worms parasitize grasshoppers and crickets, cause their hosts to enter water and thereby facilitate the release of new adult parasites, often resulting in the death of hosts (11). So-called zombie ants are compelled by infection with the parasitic fungus *Ophiocordyceps unilateralis* to leave their homes on the forest floor and attach themselves to the veins of elevated leaves; large fruiting fungal bodies grow from the ants’ heads and rupture, killing the ants and releasing fungal spores across the forest floor to start the cycle again (12). Increasingly, evidence suggests that parasite manipulation of hosts is not a new phenomenon – ant fossils provide evidence that fungal and helminth manipulation strategies were already well established 30-50 million years ago (13, 14).

Under trophically transmitted parasite systems, parasites are transmitted when hosts are preyed upon, and behavioral manipulation of hosts is therefore frequently aimed at increasing host susceptibility to predation (15). Fluke-infected California killfish display atypical swimming behavior that attracts bird predators who are required to complete the parasites’ lifecycle (16). Ants infected with the roundworm *Myrmeconema neotropicum* turn from black to bright red, then situate themselves amongst red berry clusters and raise their abdomens, thereby attracting frugivorous birds that inadvertently consume infected ants and serve as the parasite’s final host (17).

One of the best known examples of behavioral manipulation aimed at increasing predation by definitive hosts comes from the protozoan *Toxoplasma gondii*. The only known definitive hosts of *T. gondii* – hosts in which sexual reproduction is possible - are members of the Felidae family (domestic cats and their relatives). Infected cats shed enormous numbers of unsporulated oocysts in their feces for 1-3 weeks. Shed oocysts sporulate within 1-5 days, becoming infective to the next host. Oocysts persist in the environment and on surfaces for extended periods of time, where they can be accidentally ingested by both felids and intermediate hosts (18). Intermediate hosts sustain an asexual reproduction component of the *T. gondii* lifecycle, which can occur within all warm-blooded animals, including humans (19). To complete the life cycle, cats must ingest infected intermediate hosts. While directly testing the effect of *T. gondii* infection in human neural systems is prohibitively invasive, this work has been done in rodents, and sheds some light on post-infection processes in intermediate hosts. In intermediate hosts, *T. gondii* replicates rapidly in tissues until suppressed by the host’s immune response. Some parasites differentiate into slowly replicating bradyzoite forms contained within tissues cysts, most commonly in the brain and muscles, resulting in life-long infection (20). Behavioral experiments have shown that infected rodents lose their innate fear of cats, and in some cases show a fatal preference for cat-related olfactory signals (21-23). For a trophically transmitted organism, the fitness benefit to the parasite is clear: increasing the likelihood of transmission to the feline definitive host by increasing the likelihood of predation on infected intermediate hosts.

While a major aim of infection of intermediate hosts by *T. gondii* is the behavioral manipulation of prey animals, ingested oocysts infect many non-prey organisms and undergo similar processes after ingestion (19). It has been estimated that 30-50% of the global human population may be chronically infected with *T. gondii*, and a recent study estimated that up to 84% of France’s population is infected (24, 25). Upon infection, humans sometimes exhibit mild, flu-like symptoms in the first weeks following exposure, after which no clinical symptoms are regularly associated with infection in immunocompetent adults (26). Despite the absence of clinical pathology in chronically infected humans, there is some evidence linking *T. gondii* infection to variation in human behavior. Research on the effects of human infection has uncovered correlations between several behavioral and clinical outcomes and *T. gondii* infection status (27). *T. gondii* infection has been found to influence personality profiles (28, 29), reaction time of infected subjects (30), increased height and aggression in males (31), changes to female perceptions of masculinity (32), an increased risk of traffic accidents (33), and the risk of suicide (34, 35). Clinically, at least 40 studies have found an increased prevalence of chronic, latent *T. gondii* infection among schizophrenic patients (36), and amongst mothers of children with Down syndrome (37). It is unknown whether these mixed bags of symptoms and syndromes relate to mechanisms aimed at manipulating prey animals (38); indeed, they may be mere immunological side effects of chronic infection in humans (39, 40), or represent spurious conclusions drawn from meaningless correlations. To determine which, if any, human behaviors change as a result of *T. gondii* infection, requires understanding: i) potential neuroendocrine mechanisms by which highly specific infection-generated behaviors might arise, and ii) how these effects might be fixed through selection.

With respect to potential neuroendocrine mechanisms resulting in highly specific behaviors, infection should either i) target the specific brain regions that control relevant behaviors, or ii) be capable of neuromodulation through alterations in neurotransmitter levels (41). While generally not displaying tropism for specific brain regions in intermediate hosts, several studies have identified the brain regions affected by *T. gondii* infection in immunocompromised patients using MRI, particularly the frontal and parietal lobes, the cerebral cortex, the basal ganglia, and the cerebellum (42-45). Perhaps more compellingly, and supporting the potential for neuromodulation through alterations in neurotransmitter levels, encysted *T. gondii* bradyzoites can synthesize dopamine. One study found that total brain dopamine in chronically infected mice was elevated 114% over the level of uninfected mice, while other neurotransmitter levels remained unchanged (46). Another study found extremely high concentrations of dopamine in cyst-containing brain cells in infected mice, and *in vitro* infection induced high dopamine levels in neural cells (20). Intriguingly, the dysregulation of dopamine is associated with many of the human neurological disorders that are correlated with toxoplasmosis in humans, including schizophrenia (47), personality disorders (48), suicide risk (49), Tourette’s syndrome (50), autism spectrum disorders (51), and bipolar disorder (52). While not definitive, the preferential localization of *T. gondii* in the central nervous system, coupled with its ability to modify dopamine signaling in host cells, provides a reasonable potential mechanism by which chronic, latent, *T. gondii* infection might result in consistent behavioral changes in intermediate hosts.

With respect to how *T. gondii*-induced behavioral changes in humans might become fixed through selection, it is necessary to understand human-cat interactions. While humans are not regularly felid prey, the domestication of cats is a mysterious and unprecedented phenomenon. Domestic cats are one of the world’s most popular pets, however, the origin and function of cat domestication is murky, and remains an enduring source of debate (53-55). Molecular and archaeological evidence indicates that cat domestication occurred ∼10,000 years ago in the near east (53, 56, 57), at the same time as the development of human sedentism and agricultural economies (53, 58). The prevailing current hypothesis is that cats essentially domesticated themselves, in a process whereby some wildcats exploited newly developed anthropogenic environments, were tolerated by people for their utility in controlling grain pest populations (e.g. rodents), and over time human-adjacent cats and wildcats slowly diverged (59, 60).

While the utility of cats as commensal organisms under agrarian sedentism is a widely agreed-upon narrative, it is highly problematic for several reasons. First, the domestication of an apex predator is otherwise unknown in human history. Second, all felids are obligate carnivores, and unable to digest anything beyond animal protein (61), meaning regardless of their roles in pest control they would have directly competed with humans for highly valued foods. Third, despite cats’ ability to perform pest control activities, cats do not perform tasks as directed, and their actual utility as “mousers” is debatable - dogs and ferrets would be more suitable and trainable candidates (60). Fourth, unlike dogs who are socially living animals and can form strong bonds with humans (62), cats are solitary and territorial, making them more attached to places than to people. Finally, unlike all other known domesticated animals, cats are not regularly exploited for labor, meat, fur, skin, or milk. Taken together, cats are imperfect candidates for domestication, even given their tenuous utility as freelance mousers.

Given the close relationship between humans and domestic cats, it is feasible that parasite-mediated neuroendocrine mechanisms that increase human tolerance of cats would be selected for in order to favor *T. gondii* fitness through increased availability of suitable feline hosts. Here, we assess the hypothesis that *T. gondii* can change human behavior to favor the parasite by changing human behaviors that regulate the fitness of the parasite’s definitive host - cats. Specifically, we assess the possibility that human behavioral changes resulting from chronic, latent *T. gondii* infection lead to measurable changes in affection and actions towards cats that function to increase domestic cat fitness. We assessed *T. gondii* infection status in 68 participants, and assessed their attitudes towards cats in three ways: i) surveys, ii) participant behavior in the presence of domestic cats, and iii) participant oxytocin levels before and after interactions with cats to evaluate dopaminergic changes.

## Methods

### Data Collection

We assessed the effect of *T. gondii* infection on attitudes and behaviors towards cats in 68 healthy adult volunteer participants without animal allergies from the Gainesville/University of Florida community. Volunteers were scheduled for participation following poster-based recruitment efforts. Upon arrival, participants were first consented and then asked to provide the first of two 3 mL saliva samples via passive drool, which was collected into 15 mL tubes via straw and stored at -20°C within 5 minutes of collection. After the first saliva collection, participants were led to an adjoining room containing one desk, one chair, one desktop computer, and two bonded, friendly, disease-free (assessed through serology at the University of Florida Veterinary Hospital) indoor cats (Cocoa and Sundae) belonging to study staff. Cat toys, hiding boxes, litter boxes, water, and treats were always available, and both cats were free to move around the room at all times. All research and serological testing involving cats was approved by the Institutional Animal Care and Use Committee at the University of Florida (IACUC Protocol #202200000627).

Participants were deceptively told that a study staff member had to bring their pet cats to work that day, and they were asked to enter the room containing the study cats to complete a survey. After participants entered the cat room, study staff started a timer, and participants were told there were issues with the cat room computer, and they would have to be transferred to another room. Participants were then left alone in the room with the cats for a minimum of five minutes, while study staff pretended to get the other room ready. During this five-minute period, participants were filmed without their knowledge using a hidden camera installed in the ceiling. After participants were transferred to another room, they were asked to complete a ∼20-minute-long behavioral survey on a desktop computer about their attitudes and feelings towards cats, and domestic animals generally (Supp. Mat). Precisely 30 minutes after their initial exposure to the cats, participants were asked to provide a second 3 mL saliva sample, which was immediately stored at -20°C. After completion of the survey and collection of the second saliva sample, participants were fully debriefed on the deception (the true reason for the presence of cats, and that their behavior in the waiting room was filmed). After debriefing, participants were asked to provide complete informed consent, and if they did, to provide a blood sample to test for their *T. gondii* infection status. Whole blood draws were completed by an on-site phlebotomist who collected, by venipuncture, 3-5 mL of blood into a serum tube. Participants were then given the option of receiving the results of their *T. gondii* test. All human research, including deception, was approved by the Institutional Review Board for research with human subjects at the University of Florida (IRB #202101688).

### Behavioral Analysis

Five-minute-long videos of all participants while they were in the cat room were coded according to an ethogram (Supp Mat). For each video we recorded the duration of each behavior, and whether interaction behaviors were cat-initiated or participant-initiated. We then summed the duration of each behavior and divided it by the five-minute duration of each video. In total, we coded behaviors in the videos of 68 participants.

### Survey Measures and Analysis

Participants completed a 20-minute survey using Qualtrics on a desktop computer in the research space during the study assessment. The following survey measures were collected from each participant:

Sociodemographic characteristics: sex assigned at birth, gender identity, age, family income, and whether they receive financial aid to attend college.

Interactions with cats: how often they interact with cats, whether they own a cat, how many cats owned, where they obtained their cat, whether their cat has access to indoors, outdoors, or both, whether they grew up with a cat, during what ages they had a cat growing up, whether their cat while growing up had access to indoors, outdoors, or both, whether they’ve ever been injured by a cat, and whether they had ever been diagnosed with a cat-related disease.

Attitudes and affinity toward cats were assessed using two adapted scales. The first was adapted from existing studies (63) and asked the respondent to rate, on a scale of 1-10, the extent to which they agree with each statement: 1. I am a cat person; 2. I am a dog person; 3. I like cats and dogs about equally; 4. I do not like cats or dogs. The second measure was adapted from the Brief Attitudes Toward Animals Scale for Children (BATASC) (64), which was amended to specifically refer to cats in each item. The twelve-item measure included items such as “I love taking care of cats” and “playing with a cat is fun,” which participants rated on a 5-point agreement Likert scale. We summed the score for each participant, and then compared resulting scores between *T. gondii*-positive and *T. gondii*-negative participants.

Compassion toward animals: participants completed the Identification with Animals Measure (65), a 31-item measure including items such as “I feel strong ties to other animals” and “it is pleasant to be an animal.” Participants rated their agreement with each item on a 7-point Likert scale. This survey was designed to identify three dimensions by which humans identify with non-human animals: solidarity with animals, human-animal similarity, and animal pride (65). Solidarity with animals is defined by feeling connected to other animals and is associated with more contact with animals (i.e., pets) and a greater desire to help animals and to engage in collective actions on their behalf, even if this implies withdrawing privileges to humans. Human–animal similarity is defined by the perception that animals share similarities with humans; this dimension is associated with increased moral concern for their welfare and a greater attribution of typically human traits to other animals. Finally, animal pride is defined by a direct recognition and positive endorsement of the social category that includes all animals. It is associated with viewing humans as more animal-like, and with more competitive and instrumental intergroup relations.

The health of participants was measured by a series of questions related to chronic disease, other major medical conditions, environmental toxin exposure, occurrence of a fever over the past two years, antibiotics prescription in the past two years, frequency of illness during childhood, and occurrence of recent and current illnesses. Participants were also asked to report information related to their diet and lifestyle: tobacco use, physical activity, exercise, and whether they consider themselves to have a balanced diet.

To assess factors that may impact the measurement of hormones, participants who identified themselves as people who menstruate were asked to complete information regarding their use of hormonal contraceptives, as well as information about their menstrual cycle.

Personality dimensions were assessed using the HEXACO-PI-R, a 60-item personality inventory that measures the factors of Honesty-Humility, Emotionality, Extraversion, Agreeableness, Conscientiousness, and Openness to Experience (66). Each factor is comprised of several facet scores, which are obtained by summing or reverse-summing participant responses. Because of the lack of statistical power in our sample, personality facets were excluded from analyses and results – however, in a larger sample, they can be useful, particularly in assessing the facet of fearfulness that contributes to the Openness to Experience factor, given the frequent association in non-human animals of *T. gondii* infection and reduced neophobia (21, 67).

### Serological analysis of T. gondii infection status

Whole blood collected by the phlebotomist was left to clot for ∼30 minutes at room temperature, after which it was centrifuged at 1,000 RPM for 10 minutes to separate serum. After centrifugation, the serum was pipetted into a 2 mL cryotube and immediately stored at -80°C. Prior to antibody testing, serum samples were thawed at room temperature and briefly vortexed to homogenize them. Sera were tested for *T. gondii*-specific IgG antibodies using the DRG Toxoplasma gondii IgG ELISA kit (DRG International, Inc., Springfield, NJ) following the manufacturer’s instructions. The kit included positive and negative controls as well as three standards to ensure the validity of each test run and to allow for quantitation of antibody levels. All controls, standards, and samples were tested in duplicate, and wash steps were performed manually. Absorbance was measured at 450□nm with a reference wavelength of 630□nm using a BioTek 800 TS microplate reader (BioTek Instruments, Winooski, VT). Optical density (OD) values from duplicate wells were averaged to arrive at a mean OD. Results of each ELISA run were validated and interpreted according to the manufacturer’s guidelines, providing both qualitative and quantitative results. IgG concentrations (IU/mL) for each sample were interpolated based on a 4-parameter logistic curve generated using Prism software version 10.3.1 (GraphPad Software, Boston, MA).

### Oxytocin analysis

We measured peripheral oxytocin concentrations in participants’ saliva samples using a commercial enzyme immunoassay kit manufactured by Arbor Assays (Ann Arbor, MI). All oxytocin assays were conducted at the Primate Behavior Lab at the University of Michigan. After thawing, we centrifuged saliva samples at 1500 x g for 15 minutes and withdrew the clear supernatant for use in assays. In a preliminary analysis on a subset of our samples, we noted i) that sample dilution or concentration was unnecessary, as 1:1 samples typically yielded concentrations in the middle of the kit’s standard curve; and ii) when comparing results from extracted versus unextracted samples, the former exhibited unsatisfactory performance (comparatively high CVs and poorer fits to standard curves). The choice of whether or not to run an extraction step prior to oxytocin assays is a matter of debate, with no clear consensus on the superiority of one choice over the other (68). Additionally, Arbor Assay’s own materials report satisfactory parallelism and spike recovery from either method. Given these considerations, we report results from unextracted assays for our full set of samples. Intra-assay CVs of duplicate measurements averaged 11.54%. The inter-assay CV of controls across assay plates was 7.96%. Within participants, time 1 and time 2 oxytocin measurements were strongly correlated (*r* = 0.64).

## Results

Based on IgG antibody testing, 2 of 68 study participants were *T. gondii*-positive. Due to the low number of *T. gondii*-positive individuals in the study, only descriptive statistics were performed.

In total, toxoplasmosis-positive participants spent more time actively engaging with study cats (87.17% vs. 75.54%), and less time ignoring or not engaging with study cats (12.83% vs 16.47%) relative to toxoplasmosis-negative individuals (**Figure 1**). With respect to specific participant-initiated behaviors, toxoplasmosis-positive individuals (N=2) spent a greater proportion of total interaction time engaged in the following behaviors relative to toxoplasmosis-negative individuals (N=66): photographing cats, playing with cats using toys, observing cats, holding cats, and other engagement behaviors. Toxoplasmosis-negative participants spent a greater proportion of total interaction time in the following behaviors relative to toxoplasmosis-positive individuals: Seeking out study cats, actively and passively petting study cats, playing with cat toys, reaching out/presenting hands to cats, offering cat treats, and vocalizing to cats (**Figure 2**).

**Figure 1.**
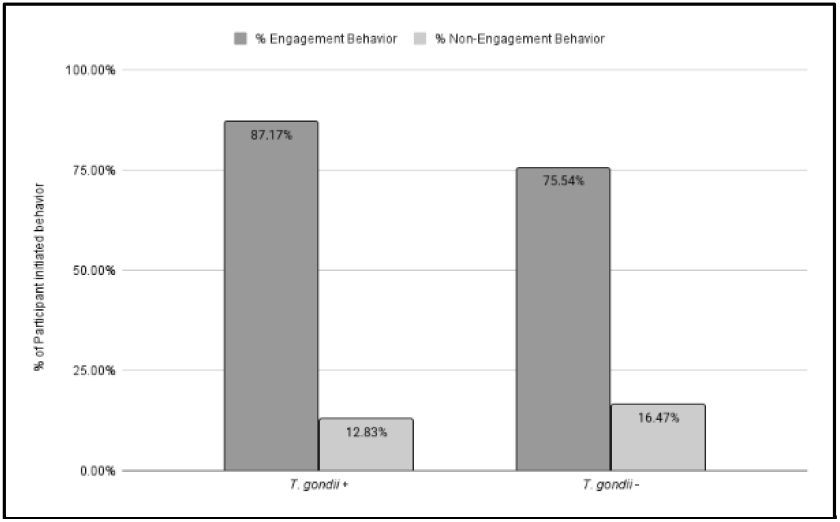
Percentage time spent in participant-initiated engagement and in non-engagement with study cats by toxoplasmosis-positive (N=2) and toxoplasmosis-negative (N=66) participants.

**Figure 2.**
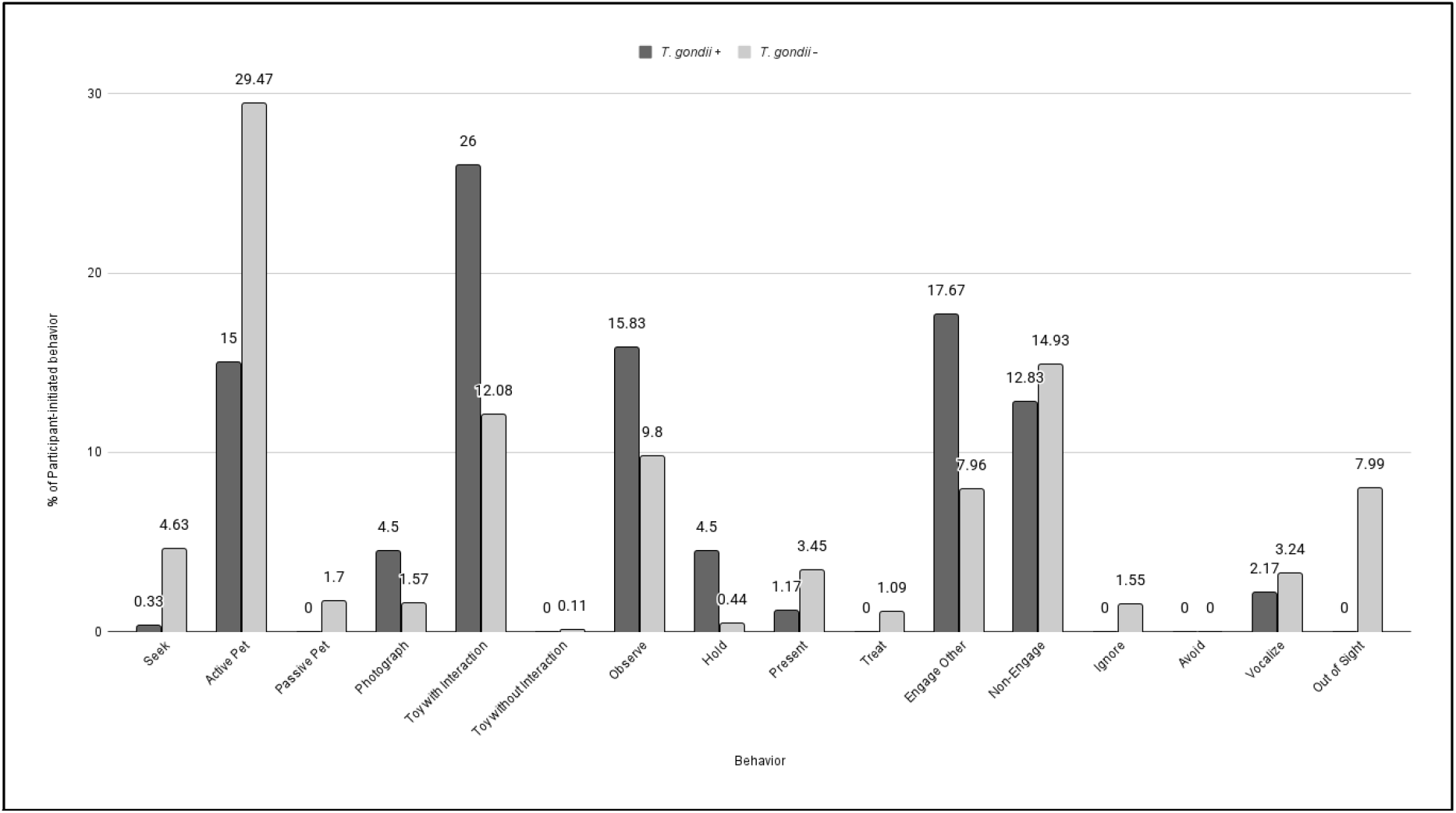
Participant-initiated behaviors as a percentage of total interaction time for toxoplasmosis-positive (N=2) and toxoplasmosis-negative (N=66) participants.

We graphed survey responses based on the maximum score achievable by summary survey statistics and survey statements, and compared responses between toxoplasmosis-positive participants (N=2) and toxoplasmosis-negative participants (N=66). First, we assessed whether infection status affected participant self-identification as a “cat person” or “dog person,” and whether they liked cats and dogs equally, or not at all. Overall, toxoplasmosis-positive participants agreed very strongly with the statement “I am a cat person” (mean=10) compared with toxoplasmosis-negative participants (mean=8.07, STD=2.23, range=2-10). Toxoplasmosis-positive participants also agreed more strongly with the statement “I like cats and dogs equally” (mean=8, range=6-10) relative to toxoplasmosis-negative participants (mean=6.5, STD=3.14) (**Figure 3**).

**Figure 3.**
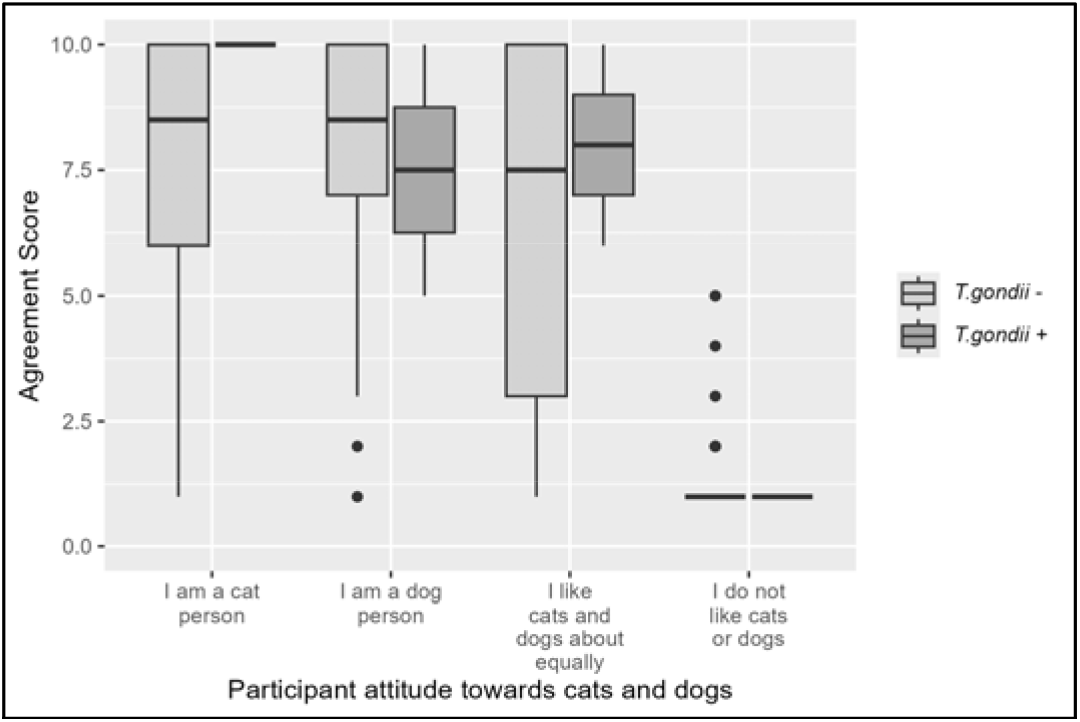
Survey responses to questions about whether cats or dogs were preferred by toxoplasmosis-positive (N=2) and –negative (N=66) participants, using a 1-10 scale, where 1 is strongly disagree and 10 is strongly agree.

The results of the modified BATASC survey similarly indicate that toxoplasmosis-positive participants report more cat-positive attitudes than toxoplasmosis-negative participants (**Figure 4**). In this survey participants were asked 12 questions which they rated on a 5-point Likert scale. Toxoplasmosis-positive participants self-reported extremely positive attitudes towards cats (mean=59, range=58-60, N=2) relative to toxoplasmosis-negative participants (mean=54.59, STD=5.04, range=38-60, N=66).

**Figure 4.**
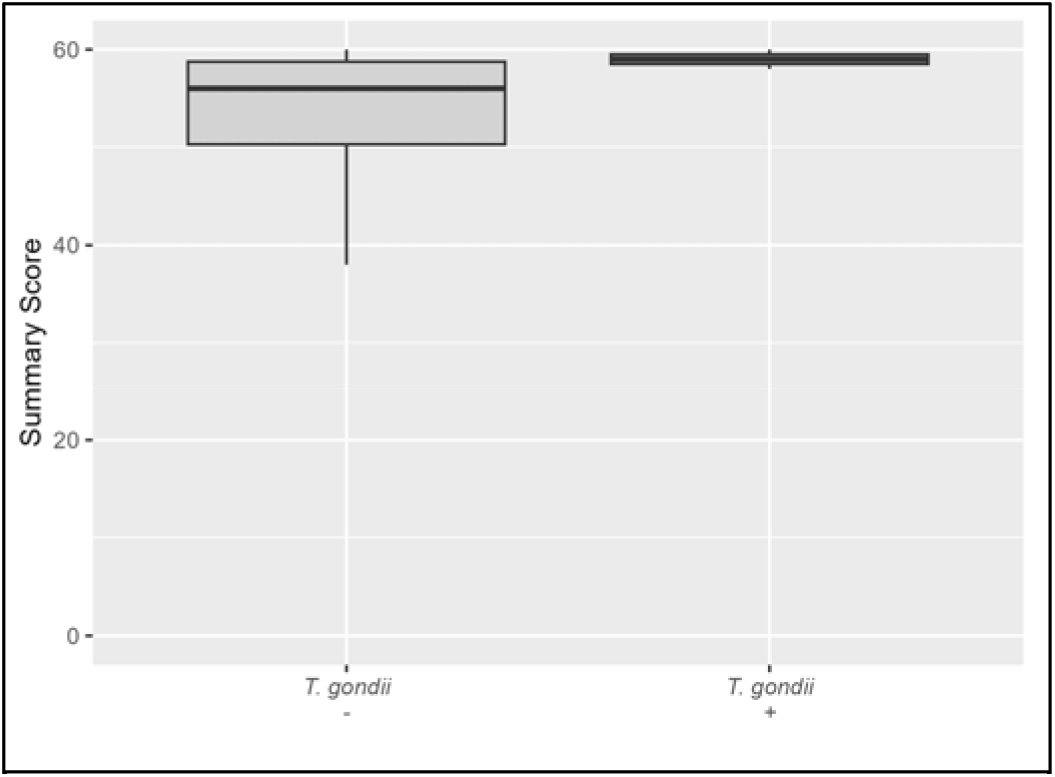
Results of modified BATASC survey, a 12-item measure to assess attitudes towards cats.

To further assess whether affection for cats is cat-specific, or part of a generalized behavioral phenotype of compassion towards animals, we employed the Identification with Animals Measure, a 31-item measure that scores three major components of human compassion for animals in general (and not cats specifically) (65) (**Figure 5**). While the low sample number precludes statistical analysis, the results indicate that toxoplasmosis-positive participants – despite identifying themselves strongly as “cat people” – scored lower or within the range of toxoplasmosis-negative people. On average, toxoplasmosis-negative participants scored higher in the dimension of solidarity with animals (mean=62.6, STD=5.04, range=46-77, N=66) than toxoplasmosis-positive participants (mean=61.5, range=60-63, N=2).

**Figure 5.**
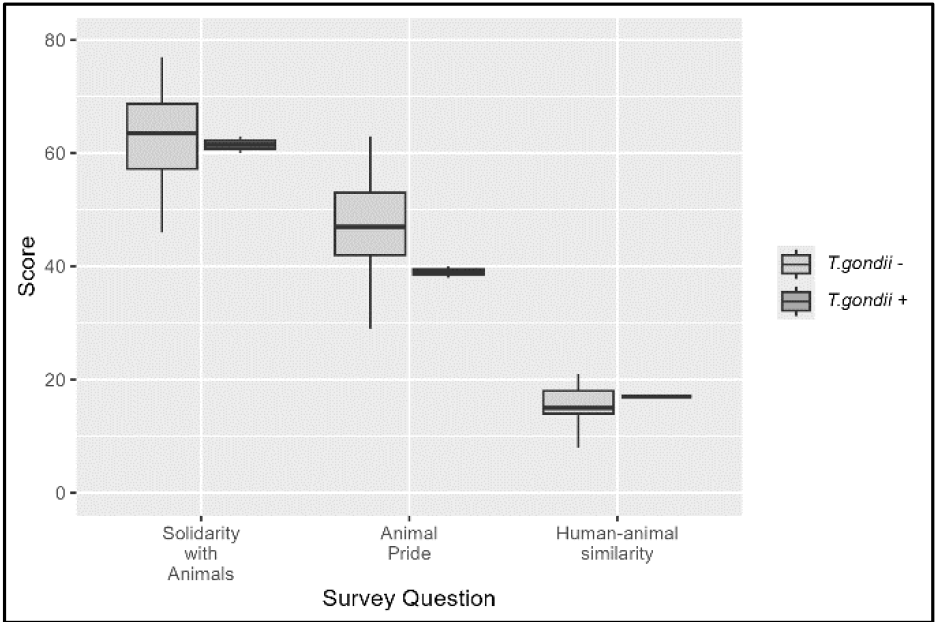
Results of Identification with Animals survey, which measures participant compassion towards animals generally across three scales.

We also assessed the difference in oxytocin concentrations before and after exposure to study cats by subtracting the post-engagement concentration from the pre-engagement concentration (**Figure 6**). Contrary to our predictions, oxytocin concentrations after exposure to cats were not higher in toxoplasmosis-positive participants than in toxoplasmosis-negative participants.

**Figure 6.**
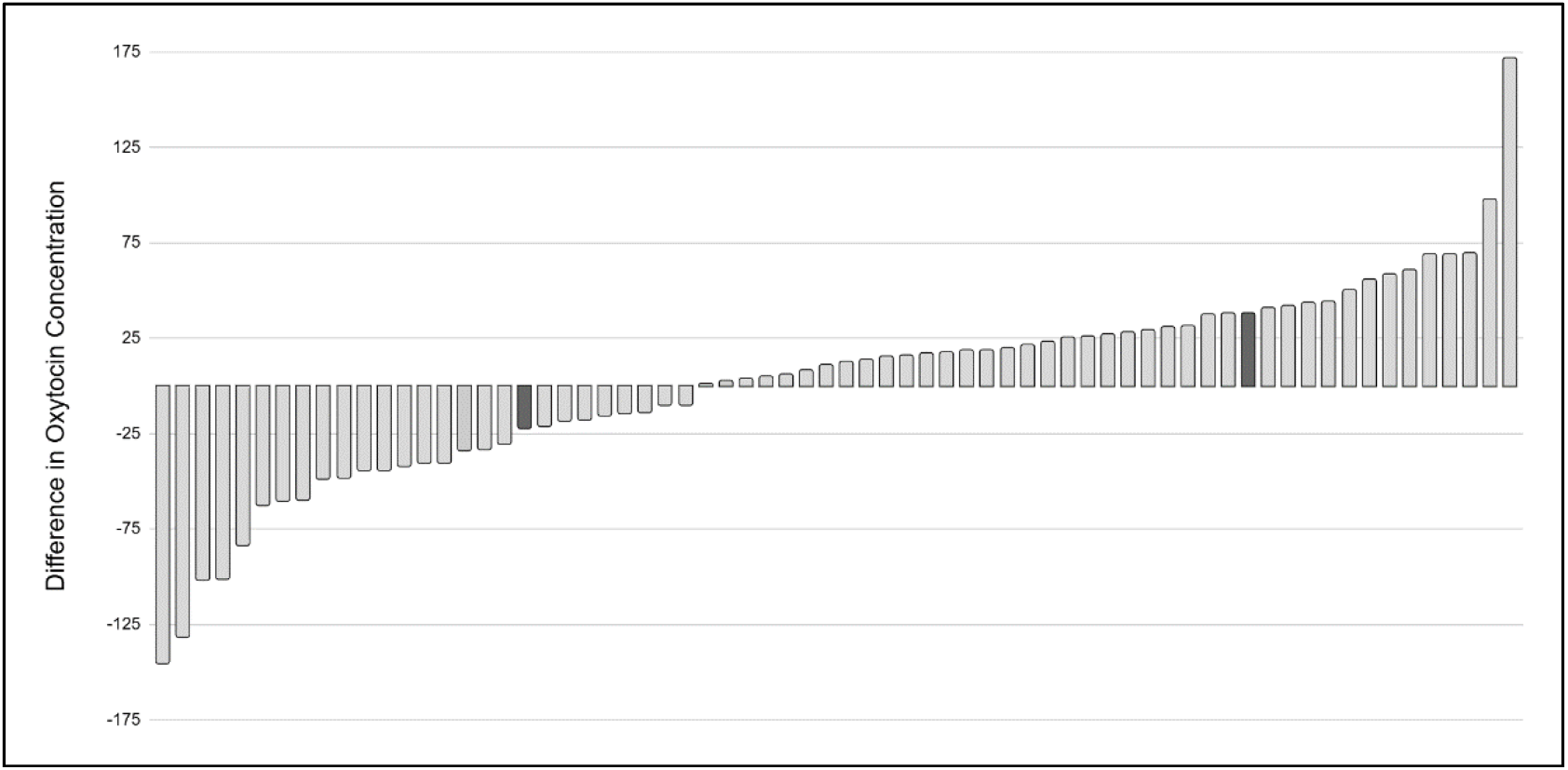
Difference in oxytocin concentrations in saliva: pre-cat exposure concentrations minus post-cat exposure concentrations, arrayed from least to most. *T. gondii-*positive participants are represented by dark bars.

## Discussion

This study sought to assess whether chronic, latent *Toxoplasma gondii* infection in humans is associated with measurable differences in behavior and attitudes toward domestic cats, the parasite’s definitive host. Building on evidence of parasite-induced behavioral manipulation in non-human hosts, particularly rodents, we tested a hypothesis that has been repeatedly proposed but rarely empirically investigated in humans: that *T. gondii* may alter human behavior in ways that ultimately favor the parasite’s fitness by promoting tolerance, attraction, or proximity to cats. Although our study included only two seropositive individuals - substantially limiting statistical inference - the pattern of results observed in this limited data set is consistent with predictions derived from the parasite manipulation hypothesis.

Across both behavioral observations and self-reported attitudes, *T. gondii*–positive participants showed stronger affection toward cats than uninfected individuals. During the covert behavioral trial, infected participants spent a higher proportion of time engaging with cats, and a lower proportion of time ignoring or avoiding them. These differences, while described qualitatively due to low sample size, point toward increased approach behaviors, similar to the reduced neophobia and cat-directed attraction documented in infected rodents (21–23). Notably, the infected participants’ behaviors tended to be exploratory, affiliative, or curiosity-driven (e.g., photographing, holding, observing), consistent with heightened incentive salience toward cat-related stimuli.

Survey results mirrored these behavioral trends. *T. gondii*–positive individuals strongly identified as “cat people,” expressed more positive attitudes toward cats, and scored near the maximum on the modified BATASC scale. The specificity of this preference is notable: infected participants did not show elevated scores on a general animal-compassion measure. This divergence suggests that the observed behavioral orientation is cat-specific rather than an artifact of broader empathy or pro-animal sentiment.

Taken together, the concordance between observed behavior and self-reported attitudes, despite an extremely small number of infected individuals, is intriguing. The directionality of these trends aligns with predictions of the manipulation hypothesis and with known neurobehavioral effects of *T. gondii* in other hosts.

Contrary to our predictions, we did not observe greater oxytocin reactivity among *T. gondii*–positive participants following cat exposure. This null pattern is difficult to interpret given the small sample size of infected participants and the ongoing debate about the validity of peripheral oxytocin measures (68). Oxytocin is known to act in part by modulating dopaminergic pathways related to social salience, reward, and motivation (69). Thus, even in the absence of measurable salivary changes, alterations in dopaminergic signaling could still provide a plausible mechanism for increased approach behaviors toward cats.

Experimental work in rodents and cell cultures shows that bradyzoite cysts can synthesize dopamine (20, 47), potentially elevating local dopamine levels in host neural cells. Dopamine confers motivational salience—shifting stimuli toward either increased attractiveness or reduced aversion. This is consistent with the dual potential mechanisms through which *T. gondii* could influence human behavior: i) incentivizing approach toward cats, or ii) diminishing avoidance or negative responses to cats. The behavioral pattern observed in the present study – higher engagement, willingness to interact physically, and increased interest – may reflect increased incentive salience of cat stimuli.

A critical challenge, and one emphasized by prior work, is distinguishing parasite-driven behavioral manipulation from baseline personality traits that predispose individuals to become infected (or to keep cats). In other words: do cat-loving people become infected, or does infection increase affinity for cats? Although our design cannot resolve this causality problem, existing longitudinal and cross-sectional evidence provides useful context. Several large-scale studies show that behavioral differences between infected and uninfected people increase with time since infection (70), strongly suggesting behavioral effects arise after infection rather than that personality predicts exposure. Additionally, our finding that infected participants did not differ from uninfected individuals in general animal solidarity argues against a simple pre-existing empathy explanation.

Another alternative explanation is that positive attitudes toward cats may be an immunological or systemic side effect of chronic infection rather than an adaptive manipulation. This is a valid concern, and one that applies broadly across the *T. gondii*–human literature. However, from an evolutionary standpoint, what matters is not whether the mechanism is direct or a byproduct, but whether the resulting behavioral shift consistently favors parasite transmission. Behavioral consequences that enhance definitive host abundance, reduce human aversion to cats, or promote cat–human cohabitation would all increase environmental oocyst exposure and thus parasite fitness.

Testing parasite manipulation in humans is inherently difficult because humans are not typically prey animals. However, in this system, predation is not required for humans to function as intermediate hosts: humans modify landscapes, maintain cat populations, and facilitate conditions for oocyst persistence. Any behavioral changes that increase cat welfare, human–cat proximity, or human tolerance for cats could indirectly increase parasite transmission opportunities.

Domestication history further complicates interpretation. Cats are an unusual domesticate—solitary, minimally useful to humans, and poor candidates for taming based on typical domestication criteria. Several scholars have argued that the tight human–cat relationship is puzzling from a functional perspective (53–60). A parasite-mediated contribution to human tolerance or affinity for cats, even if modest, would offer a provocative complementary hypothesis.

The present study has several key limitations, primarily the extremely low number of toxoplasmosis-positive participants. Despite this limitation, the presence of consistent, directional patterns across independent measurement modalities - behavioral, attitudinal, and neuroendocrine - suggests that a larger study is warranted.

The patterns observed here motivate several key avenues for future research. Ideally, future research will include a larger sample population, perhaps by targeting populations with higher known seroprevalence of toxoplasmosis. Our hypothesis might also be tested via other noninvasive proxies for dopaminergic signaling to clarify mechanistic pathways. Furthermore, given dopamine’s role in motivational salience, future work could test whether *T. gondii* alters responses to other salient stimuli, or whether effects are cat-specific.

## Conclusion

Although based on only two seropositive individuals, our findings provide suggestive evidence that latent *T. gondii* infection may be associated with increased positive attitudes and behaviors toward domestic cats. These preliminary results are consistent with predictions of the parasite manipulation hypothesis and align with known neurobiological effects of *T. gondii* in other intermediate hosts. They also raise the possibility that even modest, subclinical behavioral shifts in humans could have ecological significance for parasite transmission. A larger-scale, well-powered study is needed to determine whether these intriguing patterns represent genuine parasite-induced modulation. Nonetheless, our findings demonstrate the feasibility of studying parasite–behavior interactions in humans and highlight an underexplored frontier in host–parasite ecology.

