## Supplemental Materials for "The effect of chronic, latent *Toxoplasma gondii* infection on human behavior: Testing the parasite manipulation hypothesis in humans"

**Supplementary Material**

**Complete Survey**

**Part 1: Demographics & Background**

1. What sex were you assigned at birth? (The next question will ask about your current gender identity)
   a. Male
   b. Female
   c. Prefer not to answer
2. What is your current gender identity?
   a. Man
   b. Woman
   c. Transgender
   d. Gender non-conforming/Non-binary
   e. A gender not listed here: ____________ (*free text response)*
   f. Prefer not to answer
3. What is your age? _______ years
4. In which of the following groups did your total family income, from all sources, before taxes, fall last year?
   a. $0-9,999
   b. 10,000-24,999
   c. 25,000-49,999
   d. 50,000-74,999
   e. 75,000-99,999
   f. 100,000-149,999
   g. 150,000 and greater
   h. Prefer not to say
5. If you are currently attending college/university, do you receive any kind of financial aid to attend?
   a. Yes
   b. No
   c. Prefer not to say

6. How often do you interact with cats?

a. Never

b. Once a month

c. Once a week

d. Once every few days

e. Everyday

7. Do you own a cat?

a. Yes

b. No *(if no, skip to #11)*

8. How many cats do you own? __________ # cats

9. Where did you obtain your cat/cats?

a)	Adoption from shelter or rescue

b)	Purchase from store

c)	Found as stray

d)	Received as gift

e)	Adopted/rehomed from friend/family member/acquaintance

f)	Other (please describe): ____________

10. Is your cat:

a. indoor only
b. outdoor only
c. Indoor and outdoor

11. Did you grow up with a cat or cats?

a. Yes
b. No *(If no, skip to #14)*

12. If yes, please circle all ages at which you/your family had a cat:

a. 0-5

b. 6-10

c. 11-15

d. 16-18

13. Growing up, were your cats:

a. Indoor only
b. Outdoor only
c. Indoor and outdoor.

14. Has a cat ever broken your skin by biting or scratching you?

a. Yes

b. No

15. Have you ever been diagnosed with a cat-related disease?

a. Yes

b. No (*if no, skip to next section)*

16. What cat-related disease have you been diagnosed with? _____________

**Part 2: Closed-Ended Questions**

Please rate, on a scale of 1-10, how much you agree with the following statements:

1. I am a cat person _____ (# 1-10)

2. I am a dog person ______ (# 1-10)

3. I like cats and dogs about equally ____ (# 1-10)

4. I do not like cats or dogs _____ (# 1-10)

**Attitudes toward cats scale**

Please indicate how much you agree or disagree with each statement below by marking an “X” in the box:

|  | Strongly agree | Agree | Neither agree nor disagree | Disagree | Strongly disagree |
| --- | --- | --- | --- | --- | --- |
| Playing with a cat is fun |  |  |  |  |  |
| Touching a cat scares, fears, or upsets me |  |  |  |  |  |
| It is strange to talk to a cat |  |  |  |  |  |
| A pet cat can make its owner happy |  |  |  |  |  |
| Having a pet cat at home is a bad idea |  |  |  |  |  |
| A cat can be a friend |  |  |  |  |  |
| When I see a cat that is hurt, I feel upset |  |  |  |  |  |
| When I know that a cat is lost, I feel upset |  |  |  |  |  |
| When someone mistreats a cat, I feel upset |  |  |  |  |  |
| A pet cat is more trouble than it is worth |  |  |  |  |  |
| I love taking care of cats |  |  |  |  |  |
| When someone annoys or frightens a cat, I feel upset |  |  |  |  |  |

**Identification with Animals Measure**

The term ‘animal’ is often reserved for non-human animals, but in fact it is a category that both humans and other animals belong to. Keeping this broader and more inclusive definition of ‘animal’ in mind, please respond to the following questions.

Please indicate how much you agree or disagree with each statement by marking an “X” in the box:

|  | Very strongly agree | Strongly agree | Agree | Neither agree nor disagree | Disagree | Strongly disagree | Very strongly disagree |
| --- | --- | --- | --- | --- | --- | --- | --- |
| I feel solidarity with animals |  |  |  |  |  |  |  |
| I feel committed to animals |  |  |  |  |  |  |  |
| I feel a strong connection to other animals |  |  |  |  |  |  |  |
| I am glad to be an animal |  |  |  |  |  |  |  |
| I think animals have a lot to be proud of |  |  |  |  |  |  |  |
| It is pleasant to be an animal |  |  |  |  |  |  |  |
| Being an animal gives me a good feeling |  |  |  |  |  |  |  |
| I have a lot in common with the average animal |  |  |  |  |  |  |  |
| I am similar to the average animal |  |  |  |  |  |  |  |
| Animals, including human animals, have a lot in common with each other |  |  |  |  |  |  |  |
| Animals, including human animals, are very similar to each other |  |  |  |  |  |  |  |
| Being an animal just feels natural to me |  |  |  |  |  |  |  |
| Animals are an important group to me |  |  |  |  |  |  |  |
| It would be a substantial risk to rescue an animal in trouble |  |  |  |  |  |  |  |
| In a time of personal need, I can rely on animals |  |  |  |  |  |  |  |
| Animals need to stick together |  |  |  |  |  |  |  |
| The group of animals is more like a collection of separate individuals than a whole |  |  |  |  |  |  |  |
| When I am with animals, I feel like we are separate individuals |  |  |  |  |  |  |  |
| I feel a kinship of sorts with other animals |  |  |  |  |  |  |  |
| I am proud to be an animal |  |  |  |  |  |  |  |
| It is good to be an animal |  |  |  |  |  |  |  |
| I am a typical animal |  |  |  |  |  |  |  |
| I don’t act like a typical animal |  |  |  |  |  |  |  |
| I have a number of qualities typical of animals |  |  |  |  |  |  |  |
| I often exhibit my positive feelings about animals |  |  |  |  |  |  |  |
| Animals are a lot alike in many respects |  |  |  |  |  |  |  |
| I feel good about animals |  |  |  |  |  |  |  |
| I am like other animals |  |  |  |  |  |  |  |
| I have a lot in common with other animals |  |  |  |  |  |  |  |
| I feel strong ties to other animals |  |  |  |  |  |  |  |
| Generally, I feel good when I think about myself as an animal |  |  |  |  |  |  |  |

**General Health and Chronic disease questions:**

Have you ever suffered or currently suffer from a chronic disease (yes/no)? ______

If so, please briefly explain your diagnosis (if known) or symptoms (if no diagnosis known):

Please check if you have ever been diagnosed with the following conditions:

_____Diabetes- Type 1

_____Diabetes- Type 2

_____Diabetes- Gestational

_____Heart disease

_____Heart defect (note type:__________________)

_____Hypertension

_____Metabolic syndrome

_____Cancer (note type: __________________)

_____Underactive thyroid (for instance, Hashimoto’s thyroiditis)

_____Overactive thyroid (for instance, Grave’s disease)

_____Multiple Sclerosis

_____Lupus

_____Rheumatoid Arthritis

_____Crohn’s disease

_____Celiac disease

_____Autoimmune disease (note type: __________________)

Have you been diagnosed with any other major medical condition (e.g., a liver disease, a kidney disease)?

____ Yes (please explain_______________________________________)

____ No

Do you work in an environment that exposes you to toxins on a regular basis? (This could be fumes from car exhaust, paint, pesticides, asbestos, radiation, mercury, etc.)

____Yes (please explain here:________________________________________)

____No

How many times in the **past two years** have you had a fever of at least 100 degrees?

____times

How many times in the **past two years** have you been prescribed antibiotics by a doctor?

____times

Relative to other students, how many days of elementary school, junior high, and high school did you miss because you were sick?

1	2	3	4	5	6	7

**Much Less	Average	Much More**

Have you been sick at all within the **last two weeks**?

____Yes, I was sick for approx. _____ days. (Type of illness:_______________)

____ No

Are you currently sick?

____Yes (Type of illness:_______________)

____No

**Diet & Lifestyle**

Do you smoke tobacco?

_____Yes, 1 to 5 cigarettes a day

_____Yes, 6 to 10 cigarettes a day

_____Yes, more than 11 cigarettes a day

_____No, I have stopped smoking (I previously smoked regularly for ______ years)

_____No, I have never smoked

Relative to other students at your university, how physically active would you rate yourself, on average?

1	2	3	4	5	6	7

**Much Less	Average	Much More**

Relative to other students at your university, how much *aerobic, stamina-building* exercise do you do, typically?

1	2	3	4	5	6	7

**Much Less	Average	Much More**

Relative to other students at your university, how much *anaerobic, strength-building* exercise do you do, typically?

1	2	3	4	5	6	7

**Much Less	Average	Much More**

Have you engaged in exercise in the last *2 hours*? (Count only exercise at least as intense and extended as jogging at a moderate pace for two miles.)

____Yes

____No

If so, what exercise? ______________

How many times have you engaged in exercise within the last *2 days*? (Count only exercise at least as intense and extended as jogging at a moderate pace for two miles.) Circle a number.

0	1	2	3	4+

Relative to other same-aged people, how balanced do you consider your diet?

1	2	3	4	5	6	7

**Very Unbalanced	Average	Very Balanced**

**Contraceptive pill and menstrual cycle questionnaire: For people who menstruate**

The following questions are for people who menstruate. Please skip to the next section if this does not apply to you.

Today’s Date _______________

Do you currently use a contraceptive pill?

___ Yes (Which brand? _________________________)

___ No

If you do not currently use the pill, have you previously?

___ Yes	When did you last use it? _________________

___ No

Do you currently use a contraceptive injection such as Depo-Provera or an implant such as Nexplanon?

___ Yes (Which brand? _________________________)

___ No, and I have never used one

___ No, but I have before; my last injection/implant was ______ (month), ____ (year)

Do you currently use an IUD, such as a copper IUD or a hormonal IUD (e.g. Mirena)?

___ Yes (Which brand? _________________________)

___ No, and I have never used one

___ No, but I have had one before; I last had it ______ (month), ____ (year)

When did your last menstrual period begin? That is, what was the first day of menstrual flow during your last period? Please state the precise date, if possible. Feel free to consult your phone’s calendar, or one we’ve provided, to identify the date if it is helpful. (If you are currently menstruating, list the date your current menstruation began.)

________________________________

What is the usual length of your menstrual cycle? (That is, what is the typical time span between the first day of menstrual blood flow one month, to the first day of menstrual blood flow the next month?) (Circle one.)

In days:	<22	22	23	24	25	26	27	28

29	30	31	32	33	34	35	36

37	38	>38

How much does the length of your cycle vary from month to month?

____ My cycle is almost always the same length month to month

____ My cycle is practically always within a day or two of the same length each month

____ My cycle is usually within a week of the same length each month

____ My cycle length is highly irregular; I never know when I’ll begin menstruating

**Personality Dimensions Questionnaire**

Please indicate how much you agree or disagree with each statement below by marking an “X” in the box:

|  | Strongly Agree | Agree | Neutral | Disagree | Strongly Disagree |
| --- | --- | --- | --- | --- | --- |
| **1. I would be quite bored by a visit to an art gallery.** |  |  |  |  |  |
| **2. I plan ahead and organize things, to avoid scrambling at the last minute.** |  |  |  |  |  |
| **3. I rarely hold a grudge, even against people who have badly wronged me.** |  |  |  |  |  |
| **4. I feel reasonably satisfied with myself overall.** |  |  |  |  |  |
| **5. I would feel afraid if I had to travel in bad weather conditions.** |  |  |  |  |  |
| **6. I wouldn't use flattery to get a raise or promotion at work, even if I thought it would succeed.** |  |  |  |  |  |
| **7. I'm interested in learning about the history and politics of other countries.** |  |  |  |  |  |
| **8. I often push myself very hard when trying to achieve a goal.** |  |  |  |  |  |
| **9. People sometimes tell me that I am too critical of others.** |  |  |  |  |  |
| **10. I rarely express my opinions in group meetings.** |  |  |  |  |  |
| **11. I sometimes can't help worrying about little things.** |  |  |  |  |  |
| **12. If I knew that I could never get caught, I would be willing to steal a million dollars.** |  |  |  |  |  |
| **13. I would enjoy creating a work of art, such as a novel, a song, or a painting.** |  |  |  |  |  |
| **14. When working on something, I don't pay much attention to small details.** |  |  |  |  |  |
| **15. People sometimes tell me that I'm too stubborn.** |  |  |  |  |  |
| **16. I prefer jobs that involve active social interaction to those that involve working alone.** |  |  |  |  |  |
| **17. When I suffer from a painful experience, I need someone to make me feel comfortable.** |  |  |  |  |  |
| **18. Having a lot of money is not especially important to me.** |  |  |  |  |  |
| **19. I think that paying attention to radical ideas is a waste of time.** |  |  |  |  |  |
| **20. I make decisions based on the feeling of the moment rather than on careful thought.** |  |  |  |  |  |
| **21. People think of me as someone who has a quick temper.** |  |  |  |  |  |
| **22. On most days, I feel cheerful and optimistic.** |  |  |  |  |  |
| **23. I feel like crying when I see other people crying.** |  |  |  |  |  |
| **24. I think that I am entitled to more respect than the average person is.** |  |  |  |  |  |
| **25. If I had the opportunity, I would like to attend a classical music concert.** |  |  |  |  |  |
| **26. When working, I sometimes have difficulties due to being disorganized.** |  |  |  |  |  |
| **27. My attitude toward people who have treated me badly is “forgive and forget”.** |  |  |  |  |  |
| **28. I feel that I am an unpopular person.** |  |  |  |  |  |
| **29. When it comes to physical danger, I am very fearful.** |  |  |  |  |  |
| **30. If I want something from someone, I will laugh at that person's worst jokes.** |  |  |  |  |  |
| **31. I’ve never really enjoyed looking through an encyclopedia.** |  |  |  |  |  |
| **32. I do only the minimum amount of work needed to get by.** |  |  |  |  |  |
| **33. I tend to be lenient in judging other people.** |  |  |  |  |  |
| **34. In social situations, I’m usually the one who makes the first move.** |  |  |  |  |  |
| **35. I worry a lot less than most people do.** |  |  |  |  |  |
| **36. I would never accept a bribe, even if it were very large.** |  |  |  |  |  |
| **37. People have often told me that I have a good imagination.** |  |  |  |  |  |
| **38. I always try to be accurate in my work, even at the expense of time.** |  |  |  |  |  |
| **39. I am usually quite flexible in my opinions when people disagree with me.** |  |  |  |  |  |
| **40. The first thing that I always do in a new place is to make friends.** |  |  |  |  |  |
| **41. I can handle difficult situations without needing emotional support from anyone else.** |  |  |  |  |  |
| **42. I would get a lot of pleasure from owning expensive luxury goods.** |  |  |  |  |  |
| **43. I like people who have unconventional views.** |  |  |  |  |  |
| **44. I make a lot of mistakes because I don’t think before I act.** |  |  |  |  |  |
| **45. Most people tend to get angry more quickly than I do.** |  |  |  |  |  |
| **46. Most people are more upbeat and dynamic than I generally am.** |  |  |  |  |  |
| **47. I feel strong emotions when someone close to me is going away for a long time.** |  |  |  |  |  |
| **48. I want people to know that I am an important person of high status.** |  |  |  |  |  |
| **49. I don’t think of myself as the artistic or creative type.** |  |  |  |  |  |
| **50. People often call me a perfectionist.** |  |  |  |  |  |
| **51. Even when people make a lot of mistakes, I rarely say anything negative.** |  |  |  |  |  |
| **52. I sometimes feel that I am a worthless person.** |  |  |  |  |  |
| **53. Even in an emergency I wouldn’t feel like panicking.** |  |  |  |  |  |
| **54. I wouldn’t pretend to like someone just to get that person to do favors for me.** |  |  |  |  |  |
| **55. I find it boring to discuss philosophy.** |  |  |  |  |  |
| **56. I prefer to do whatever comes to mind, rather than stick to a plan.** |  |  |  |  |  |
| **57. When people tell me that I’m wrong, my first reaction is to argue with them.** |  |  |  |  |  |
| **58. When I’m in a group of people, I’m often the one who speaks on behalf of the group.** |  |  |  |  |  |
| **59. I remain unemotional even in situations where most people get very sentimental.** |  |  |  |  |  |
| **60. I’d be tempted to use counterfeit money, if I were sure I could get away with it.** |  |  |  |  |  |

**Ethogram**

| **Category** | **Description** |
| --- | --- |
| Seek – **S** | Moves around room looking for cat(s); moves directly towards cat(s); tries to engage with non-responsive hidden or resting cat without directly engaging. |
| Active pet - **AP** | Touching or petting cat(s), while attention/gaze is focused on cat. |
| Passive pet - **PP** | Petting cat(s) while attention is actively focused elsewhere, e.g., phone, book |
| Photograph - **PH** | Takes a photo or video of cat(s), or sets up phone to take a photo or video. |
| Toy with Interaction - **TI** | Picks up and moves a cat toy in the direction of cat(s)/interacts with 1 or both cats with a cat toy. |
| Toy without Interaction - **TN** | Picks up a cat toy but does not use toy to approach or interact with cat(s). |
| Observe - **OB** | Participant watches or glances at cat(s) without interacting physically or vocally. |
| Hold - **HO** | Picks up or holds cat, including on lap, or in arms – *hold trumps all other behaviors*, e.g., record the behavior as HO even if while holding the cat the participant is vocalizing, petting, or otherwise engaging with cat. |
| Present - **PR** | Participant offers hand or other body part to cat(s) to sniff. |
| Treat - **TR** | Person opens treat drawer, person handles treat bag, incl shaking, and/or person offers and feeds treat. |
| Engage Other – **EO** | Interacting with cat(s) but specific behavior is unknown (i.e. can't tell precisely if it is pet, toy, present, treat, photograph, etc.) |
| Non-Engage - **NE** | Not interacting with cat(s) and cat(s) is/are not actively interacting or seeking person’s interest, e.g. sending a text, looking at pictures, reading a book |
| Ignore - **IG** | Non-engagement when cat(s) is/are actively interacting or seeking out person, e.g., cat is sniffing, rubbing on participant, approaching participant, but participant does not respond. |
| Avoid - **AV** | Participant moves away from approaching cat/cats, or pushes away cat(s). |
| Vocalize - **VO** | Calls out to cats, makes pss psss pss or tsk noise, talks to cat, sings to cat, *only note if other behaviors not present (except observe).* |
| Out of Sight - **OOS** | Participant behavior is not possible to observe. |

For each line of behavioral code that includes one of the interaction categories (e.g., pet, observe, toy with interaction, hold etc), note if the interaction was INITIATED by the cat (C), or by the participant (P).
